# Query-driven generative AI synthesizes multi-modal spatial omics from histology

**DOI:** 10.64898/2025.12.11.693669

**Authors:** Minxing Pang, Tarun Kanti Roy, Xiaodong Wu, Kai Tan

## Abstract

Spatial omics technologies offer unprecedented insights into the cellular organization of tissues; however, they are not yet scalable for routine clinical use. In contrast, images of histological staining remain the foundation of pathological diagnosis despite lacking molecular information. Bridging this gap requires computational methods that can accurately infer spatial molecular data from histology alone. Here, we introduce TissueCraftAI, a generative artificial intelligence framework that predicts multi-modal spatial omics maps directly from standard histology images using natural language prompts. To train and validate our model, we created PRISM-12M, a large-scale dataset comprising over twelve million spatially registered histology and spatial omics image patches across fourteen tissue types from humans and mice. TissueCraftAI significantly outperforms existing methods in generating realistic histology images and predicting spatial proteomics and transcriptomics data with high fidelity. We demonstrated its utility in various downstream applications, including improving cell type annotation and enhancing the accuracy of patient survival predictions across multiple cancer types. By enabling flexible, query-driven *in silico* spatial molecular analysis using routine histology images, TissueCraftAI opens up new research avenues in computational pathology.

## Introduction

The spatial arrangement of cells and molecules within tissues is essential for their normal functions and pathogenesis^1^. The advent of spatial omics technologies, such as spatial transcriptomics and highly multiplexed proteomics, has revolutionized our ability to resolve tissue cellular architecture, offering deep insights into spatial heterogeneity, cell-cell communication, and disease pathogenesis^2^. However, they are often expensive, labor-intensive, and require specialized equipment, limiting their widespread use in both basic research and clinical diagnostics. In contrast, histological analysis of tissue sections, particularly using hematoxylin and eosin (H&E) staining, has been the bedrock of anatomical pathology for over a century. Histology slides are inexpensive to prepare, routinely produced in clinical workflows worldwide, and offer a wealth of morphological information that supports numerous medical diagnoses. Despite its widespread use, H&E histology provides only a morphological snapshot and lacks the specific molecular information captured by spatial omics. This creates a critical gap: the vast archives of existing histology slides, along with the millions more produced each year, represent a massively untapped resource of latent molecular information^3–6^. Unlocking this potential requires the development of robust computational methods that can accurately predict spatial omics data from standard H&E images^7–10^, thereby bridging the gap between morphological and molecular information.

Recent advances in computational pathology have demonstrated the feasibility of predicting molecular features from histology images. These methods include contrastive learning-based frameworks, such as OmiCLIP^8^, that align H&E images with transcriptomics through retrieval-based methods, which require millions of paired image patches. Generative models, such as HistoPlexer^9^ and ROSIE^10^, which predict fixed panels of eleven and fifty protein markers, respectively, utilize conditional generative adversarial network (GAN) or convolutional neural network architectures. While these methods have shown promising results, they share critical limitations in their neural network architectural design and user accessibility. For instance, conditional GAN architectures are challenging to train and lack explicit mechanisms for preserving fine-grained spatial tissue structures during generation^9^. Additionally, these methods focus on either RNA transcript or protein expression, but not both, or exhibit limited generalizability across diverse tissue types and disease conditions. Stable diffusion-based probabilistic models have recently superseded GANs in general image synthesis due to their superior stability and mode coverage^11,12^. However, this powerful technology has not yet been adapted for spatial omics image synthesis. Another limitation of existing methods is the lack of querying ability using natural language input for intuitive molecular target specification. Instead, existing methods require users to work with rigid, pre-defined marker outputs or gene name sentences^8,9^. The development of flexible, text-based queries is essential not only for intuitive hypothesis testing but also for future integration with autonomous AI agents. By enabling natural language-based text prompts, we will pave the way for large language model (LLM)-driven systems to dynamically query molecular maps as part of complex reasoning pipelines. In summary, there remains a critical need for a unified computational framework that integrates hypothesis-driven natural language query with state-of-the-art generative modeling to enable flexible, text-guided prediction of multi-modal spatial molecular maps.

To address current knowledge gaps, we developed TissueCraftAI, a generative AI framework that leverages a text-guided latent diffusion model to translate histology images into diverse spatial omics maps. A key innovation of our approach is the use of natural language prompts, which allows users to flexibly query for any desired molecular target (e.g., “predict CD8 protein expression”) without altering the model architecture. To ensure the generated maps are spatially faithful to the input histology, our model is conditioned on the input image using a ControlNet module^13,14^, preserving the underlying tissue structure. To power the training and rigorous evaluation of this framework, we first created PRISM-12M, the largest publicly available, spatially registered, multi-modal dataset, linking H&E images with corresponding spatial proteomics and transcriptomics from the exact same tissue sections. We demonstrate that TissueCraftAI substantially outperforms current state-of-the-art models across a range of tasks, from high-fidelity histology generation to the accurate prediction of protein and RNA spatial distributions at both the cellular and subcellular levels. We further demonstrate that the spatial molecular maps synthesized by our method can be utilized in various downstream analyses, including improving the accuracy of cell type annotation and enhancing the prognostic power of clinical survival models.

## Results

### A Large-Scale, Multi-Modal Dataset for Computational Pathology

We created PRISM-12M, a large-scale dataset comprising over 12 million spatially co-registered pairs of image patches linking H&E histology with corresponding single-cell-resolution spatial omics data (Fig. 1a). This curated dataset was generated using a custom processing pipeline that performs image quality control, co-registration, and the extraction of paired patches from whole-slide images. This process ensures direct spatial correspondence between high-quality tissue morphology and molecular data (Extended Data Fig. 1).

**Figure 1.**
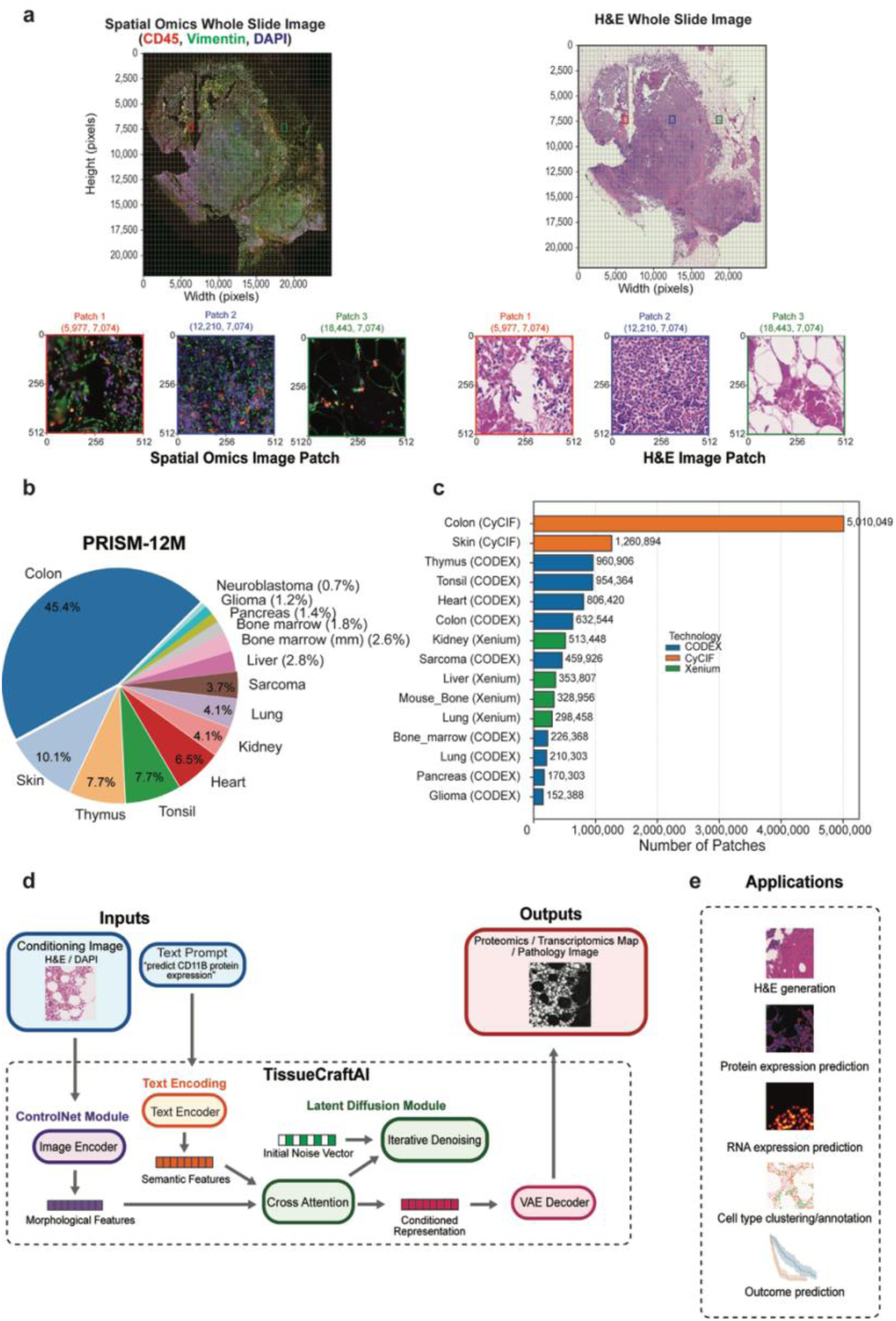
An overview of the PRISM-12M dataset and the TissueCraftAI framework. **(a)** Representative image pairs from the PRISM-12M dataset. Left, whole slide image (WSI) of spatial proteomics data overlaid with a pixel coordinate grid. The tissue was stained with anti-CD45 (red, a pan-hematopoietic marker), anti-vimentin (green, a mesenchymal/stromal cell marker) antibodies, and DAPI (blue, a nuclear stain). Right, the corresponding WSI of Hematoxylin and Eosin (H&E) staining, showing tissue morphology. The same coordinate grid is shown, demonstrating spatial alignment with the spatial omics data. Three representative regions of interest (Patches 1-3) are shown at identical pixel coordinates (Width, Height) from both images. **(b)** Pie chart showing the composition of the PRISM-12M dataset, consisting of over 12 million patches of matched H&E and spatial omics images covering 14 human and mouse tissues. Mouse tissue is indicated with a “mm” in the tissue name. The proportion of each dataset is indicated in parentheses. **(c)** Breakdown of the PRISM-12M dataset by tissue source and spatial omics data modality. CyCIF, cyclic Immunofluorescence; CODEX, CO-Detection by indEXing. (**d)** Schematic of the TissueCraftAI model. The generative model uses a dual-input, text-guided diffusion model architecture. A conditioning image (e.g., H&E or DAPI) is processed by a ControlNet image encoder to extract spatial and morphological features. Simultaneously, a user-defined text prompt specifying the molecular target is processed by a text encoder to produce semantic features. Both feature sets are integrated via a cross-attention mechanism to guide a latent diffusion module. This module iteratively refines an initial noise vector into a conditioned latent representation, which a Variational Autoencoder (VAE) decoder then translates into the final high-resolution proteomics or transcriptomics image. **(e)** Example downstream applications of TissueCraftAI, including H&E image generation, spatial protein expression prediction, spatial RNA expression prediction, cell-type clustering/annotation, and patient outcome prediction.

PRISM-12M includes fourteen different tissue types, representing both normal and pathological tissues from humans and mice (Fig. 1b). This variety of tissue types and conditions ensures that computational models developed using this dataset are generalizable across various biological and pathological contexts. Currently, PRISM-12M includes two single-cell-resolution spatial omics modalities: spatial proteomics data generated using Cyclic Immunofluorescence (CyCIF) and CO-DEtection by indeXing (CODEX) platforms, which collectively cover 166 unique proteins. Additionally, it includes spatial transcriptomics data generated using the Xenium platform, featuring 1,261 RNA species (Fig. 1c, Extended Data Fig. 2). The multi-modal nature of PRISM-12M enables the creation of unified computational frameworks that can predict protein and RNA spatial expression patterns from H&E images alone. The scale and diversity of PRISM-12M make it an invaluable resource for developing and evaluating computational methods that link tissue morphology to molecular phenotypes, addressing a critical challenge in computational pathology.

### The TissueCraftAI Framework

We introduce TissueCraftAI, a generative artificial intelligence framework for predicting spatial omics images using histology images along with natural language prompts as input. The overall architecture of TissueCraftAI is illustrated in Fig. 1d. A standard histology image, such as a H&E or a 4’,6-diamidino-2-phenylindole (DAPI) nuclear-staining slide, serves as a conditioning image from which a spatial omics image is generated. This histology image provides the essential spatial and morphological context for the tissue section. In parallel, a user-defined text prompt – such as “predict the spatial distribution of CD11B protein expression,” – specifies the molecular target for the generative task. To process the multimodal input data, TissueCraftAI employs two parallel encoders. A ControlNet neural network ^15^ encodes the conditioning image to preserve fine-grained structural details and spatial relationships from the input histology image. At the same time, a transformer-based text encoder processes the natural language query to generate a set of semantic feature vectors that capture the nuanced biological meaning of the specified molecular target.

The core of TissueCraftAI is a latent diffusion model that integrates morphological and semantic information to guide the image generation process. The generative process begins with a random noise tensor, which is iteratively refined through a series of denoising steps. At each step, a cross-attention mechanism dynamically merges the morphological features from ControlNet with the semantic features from the text encoder. This guided approach directs the denoising process towards a final latent representation that effectively reconstructs the spatial distribution of molecules. Once this compact latent representation is created, it is passed to a Variational Autoencoder (VAE) decoder, which upscales the representation and transforms it from the latent space into a final, high-resolution pixel-level image. The text-guided, diffusion-based architecture of TissueCraftAI provides significant flexibility, allowing researchers to perform hypothesis-driven queries on various molecular targets without requiring model retraining. The trained TissueCraftAI model serves as a powerful computational link between histology and spatial omics. It supports various applications, including predicting protein and RNA spatial expression, annotating cell types, and conducting association studies that link clinical variables with spatial phenotypes.

To assess the performance of TissueCraftAI, we conducted extensive benchmarking against leading image-to-image translation models across various tasks. These tasks included translating images from nucleus staining to H&E images, translating H&E images to molecular images, and annotating cell types from H&E images via predicted molecular images.

### TissueCraftAI Excels at Synthesizing Realistic Histology Images from Nuclear Staining Images

We first studied the task of translating a nuclear staining image (DAPI) to an H&E image, as these two image types are very common. We compared TissueCraftAI to two state-of-the-art image-to-image translation models, representing two distinct categories of generative models: HistoPlexe^9^, which is based on CycleGAN of Generative Adversarial Network (GAN) models, and Pix2Pix-Turb^16^, which is based on SD-Turbo of the stable diffusion models. We leveraged paired H&E and CODEX images in PRISM-12M to benchmark the performance of H&E image synthesis from DAPI images. The training and validation sets contained 4.1 million image patches, while the test set comprised 0.5 million image patches (Extended Data Fig. 1). Our quantitative assessment used two complementary metrics: the Pearson Correlation Coefficient (PCC), which measures pixel-wise correlation to quantify structural accuracy of predicted images, and the Fréchet Inception Distance (FID)^17^, which evaluates perceptual quality by comparing the statistical distribution of image features to assess biological realism.

TissueCraftAI demonstrated superior performance on both metrics, achieving the highest median PCC of 0.915, indicating a strong correlation with ground truth images (Fig. 2a). Furthermore, it obtained the lowest FID score of 138.4, significantly outperforming HistoPlexer (262.8) and Pix2Pix-Turbo (209.0), confirming its ability to generate more realistic and faithful histological images (Fig. 2b). A visual evaluation further substantiates these findings (Fig. 2c). TissueCraftAI accurately preserved fine-grained morphological details and tissue architecture, whereas images generated by the other two methods exhibited noticeable artifacts, blurring, or loss of structural integrity.

**Figure 2.**
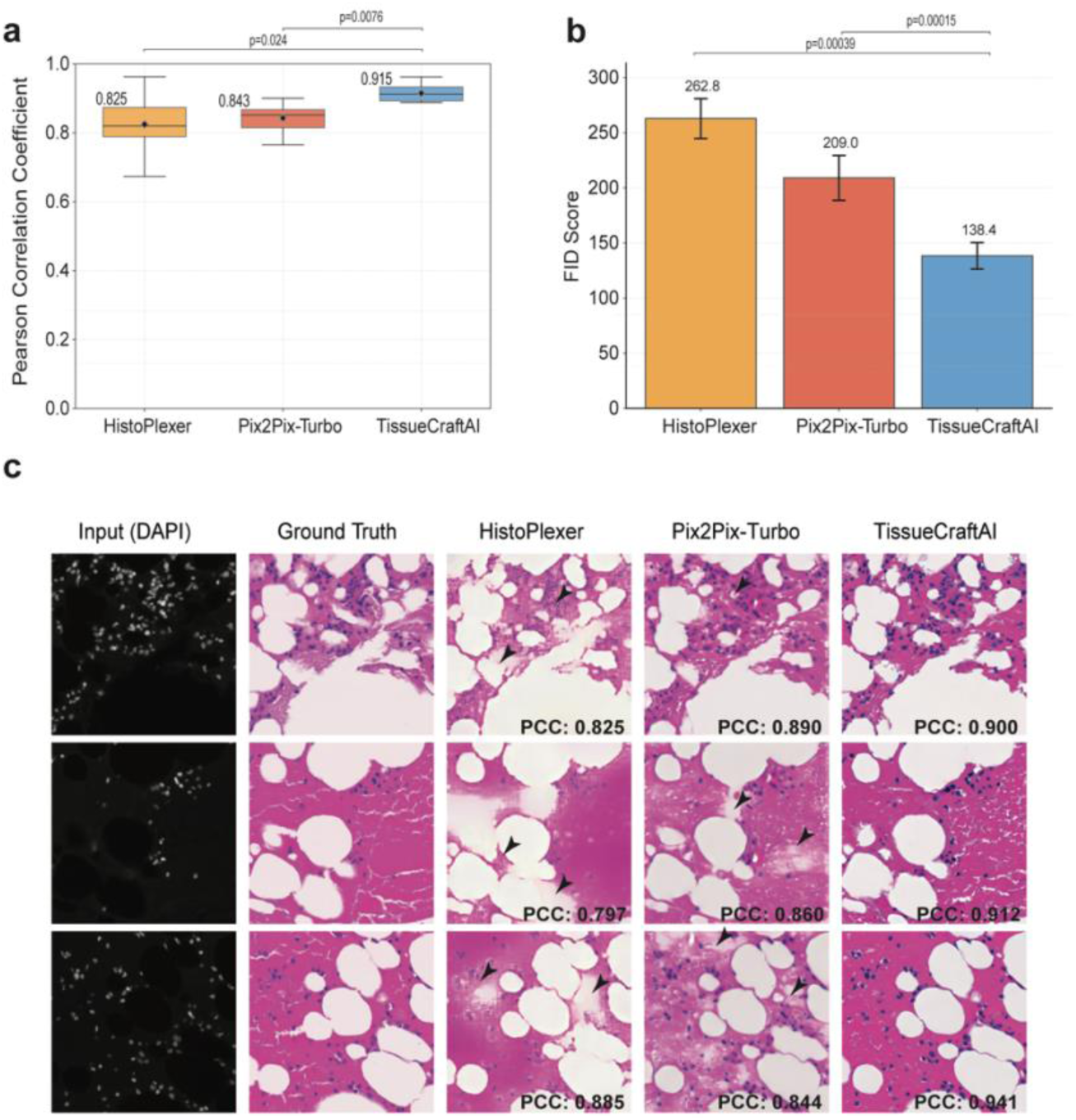
Improved histology image synthesis from nuclear staining image. **(a)** Box plot of Pearson Correlation Coefficients (PCCs) between ground truth images and images generated by HistoPlexer, Pix2Pix-Turbo, and TissueCraftAI. n=462,549 image patches. Mean values are indicated. P-values were calculated using a one-sided t-test. **(b)** Box plot of Fréchet Inception Distance (FID) scores for the three methods using all test CODEX datasets. Lower values indicate better image quality. n=8 datasets. **(c)** Representative synthetic H&E images. For three different input DAPI images, the corresponding ground truth H&E images are displayed alongside the synthesized images generated by HistoPlexer, Pix2Pix-Turbo, and TissueCraftAI. The PCCs are shown for each synthesized image. Arrowheads indicate representative image imperfections compared to the ground truth.

### TissueCraftAI Accurately Predicts Spatial Proteomics Images from H&E Images

To benchmark the performance of TissueCraftAI in synthesizing spatial proteomics images from H&E images, we compared it to HistoPlexer and Pix2Pix-Turbo. We performed a comprehensive evaluation using set-aside test datasets in PRISM-12M to assess both model accuracy and generalization capability.

First, on a human bone marrow dataset, TissueCraftAI demonstrated significantly higher predictive accuracy than both HistoPlexer and Pix2Pix-Turbo. Across a panel of key protein markers for bone marrow cell types, TissueCraftAI consistently achieved the highest PCCs, indicating a stronger agreement with the ground truth CODEX images (Fig. 3a). This global image consistency was also reflected in image fidelity. The overall quality of the generated proteomic images was superior, with TissueCraftAI achieving the best FID score of 378 compared to those of Pix2Pix-Turbo (471) and HistoPlexer (497) (Fig. 3b). A trend that held true for the set of markers across the entire PRISM-12M dataset (Extended Data Fig. 3a). Further quantifying this, we found that TissueCraftAI also yielded the highest Coefficient of Determination (R²), suggesting that its predictions account for a greater proportion of the variance in the true protein expression levels in the images (Extended Data Fig. 3b). Visual inspection corroborates these quantitative metrics. For instance, for the granulocyte marker CD15, TissueCraftAI achieved a PCC of 0.50, compared to those of HistoPlexer (0.17) and Pix2Pix-Turbo (0.15). Similarly, for the T-cell marker CD3e, TissueCraftAI’s PCC of 0.53 was nearly four times higher than those of HistoPlexer (0.11) and Pix2Pix-Turbo (0.14), demonstrating its enhanced ability to resolve fine-grained expression patterns (Fig. 3c).

**Figure 3.**
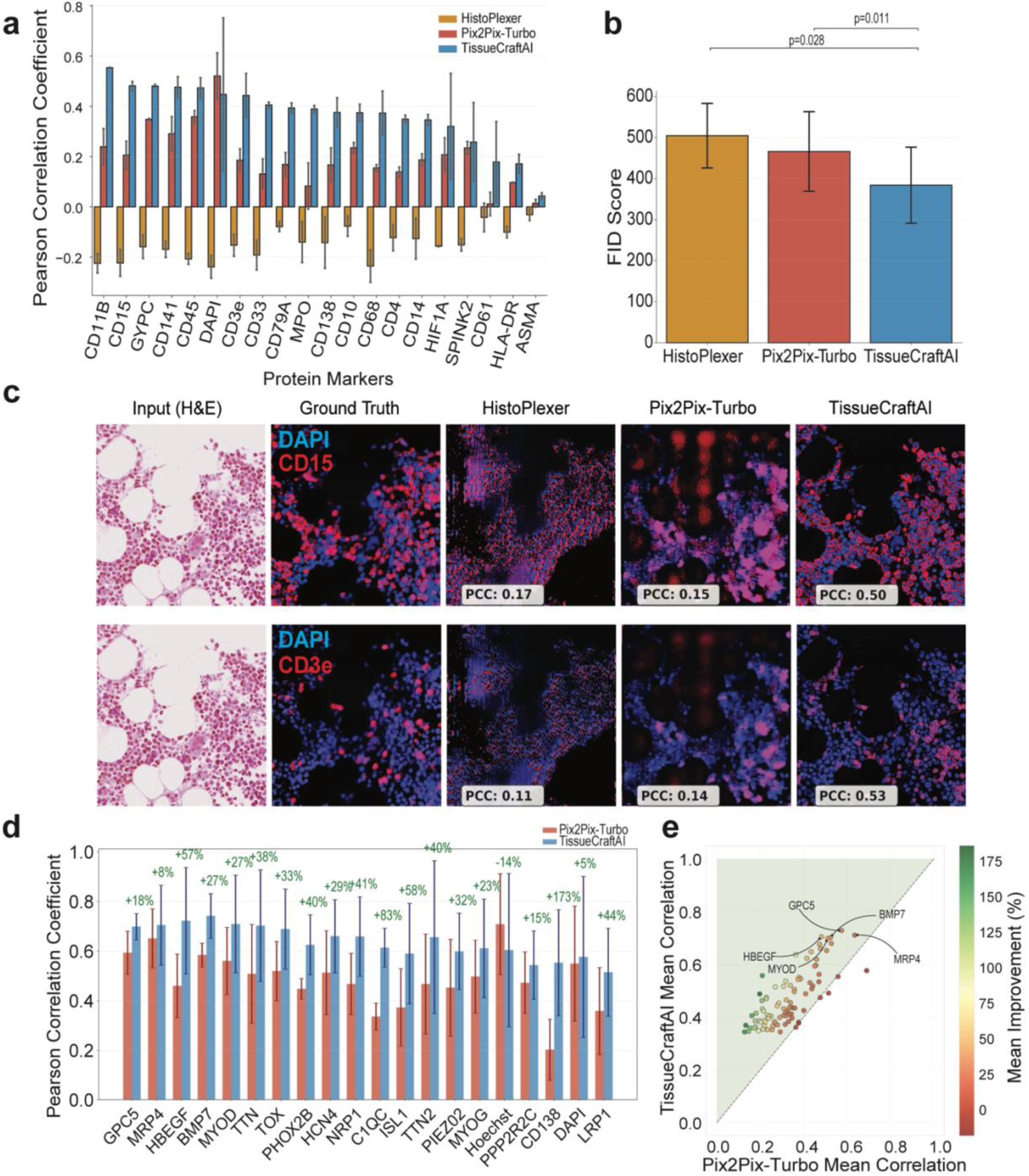
Prediction of spatial protein expression from histology images. Performance evaluation using the human bone marrow CODEX dataset. Bar plot showing the per-marker Pearson Correlation Coefficient (PCC) between the ground truth CODEX images and the synthesized images by three methods: HistoPlexer, Pix2Pix, and TissueCraftAI. Error bars represent the standard deviation. n=16,325 image patches. **(b)** Fréchet Inception Distance (FID) scores for the three methods. Error bars represent standard deviation. n=54 protein markers. **(c)** Representative examples of synthesized images. The PCC value for each synthesized image is shown. **(d)** Performance evaluation on the PRSIM-12M test set. Shown are barplots of the per-marker PCC for Pix2Pix and TissueCraftAI across all proteins in the test set. The percentage improvement of TissueCraftAI over Pix2Pix is shown above the bars. Error bars denote standard deviation. n=986,130 image patches. **(e)** Scatter plot comparing the mean correlation coefficients of TissueCraftAI (y-axis) and Pix2Pix (x-axis) for each protein. Each circle represents a protein with a corresponding mean correlation to the ground truth image. The diagonal dashed line indicates performance equivalence.

To assess its capabilities as a foundation model, we then evaluated TissueCraftAI on the entire PRISM-12M test dataset. As HistoPlexer is not a foundation model, it was not included in this more challenging benchmark. We first focused on evaluating the 20 most abundant protein markers. TissueCraftAI demonstrated superior prediction accuracy over Pix2Pix-Turbo across all markers (Fig. 3d). The performance gains were substantial for certain markers, exceeding a 50% improvement for immune markers (e.g., HBEGF, C1QC) and over 100% for the plasma cell marker CD138. To confirm that this predictive power holds broadly, we expanded the analysis to the whole test set covering 166 proteins. Our comparison shows that the vast majority of proteins fall above the equivalence line, suggesting a consistent performance gain by TissueCraftAI over Pix2Pix-Turbo (Fig. 3e). In sum, these comprehensive evaluations affirm that TissueCraftAI’s ability to learn the relationship between morphology and protein expression is robust and generalizes across a wide range of tissue types and proteins.

Finally, to assess the robustness of TissueCraftAI, we generated synthetic H&E images with common types of image artifacts and noise, including blur, lighting artifacts, smudges, and washing artifacts. TissueCraftAI was able to maintain its prediction accuracy in the presence of various types and levels of noise (Extended Data Fig. 5, and Fig. 6).

### Applications to Cellular-Level Spatial Analysis

To further evaluate the utility of TissueCraftAI for cellular-resolution spatial analysis, we benchmarked its performance on cell type annotation, a critical and challenging task in spatial omics data analysis. Starting with spatial omics images generated by TissueCraftAI, we used a standard pipeline to segment individual cells from the image, followed by quantification of molecular marker expression within each cell. The cell-by-marker expression matrices were then subjected to dimensionality reduction and clustering to group cells with similar molecular profiles. The final cell type annotation was based on known marker genes. Using H&E images as input, we compared cell type annotations generated by different methods against ground truth annotations based on real spatial proteomics data. We employed two metrics for this evaluation: the Fowlkes-Mallows Index (FMI)^18^ to measure the similarity between clusters, and the F1 score to assess the accuracy of cell type classification. The clustering-similarity-based evaluation is critical because, for many samples, cell type annotations are not available, making it challenging to resolve biologically meaningful cell clusters.

TissueCraftAI achieved the highest performance in both metrics, demonstrating its enhanced ability to predict cell types from H&E images. It achieved the highest median FMI of 0.35, significantly surpassing HistoPlexer, Pix2Pix-Turbo, and cell type annotation based solely on H&E images (Fig. 4a). Similarly, TissueCraftAI led with the highest macro F1 score of 0.38 and 0.36 using bone marrow and sarcoma H&E images, respectively (Fig. 4b, 4c). Additionally, when the cell type clustering and annotation were visualized with UMAP, the distinct clusters observed in the ground truth data were not recapitulated by HistoPlexer and Pix2Pix-Turbo, suggesting their poorer single-cell classification accuracy (Extended Data Fig. 4).

**Figure 4.**
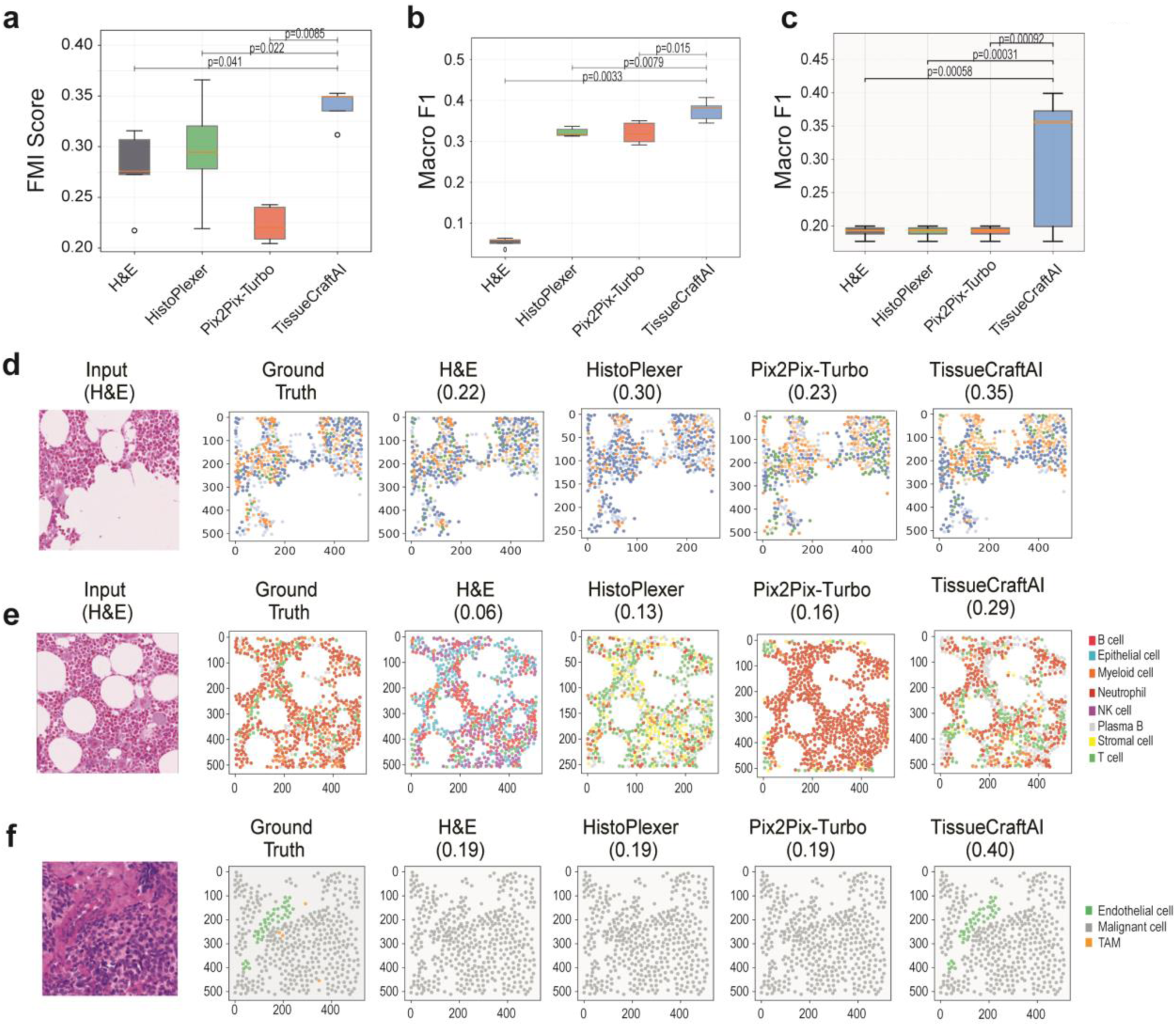
Spatially resolved cell type annotation of histology images with TissueCraftAI. **(a)** Boxplots comparing the similarity of cell type clusterings using the human bone marrow CODEX dataset. Similarity to the ground truth images is determined using the Fowlkes-Mallows Index (FMI) score for H&E images and synthetic CODEX images generated by HistoPlexer, Pix2Pix-Turbo, and TissueCraftAI. n=16,325 image patches. (**b)** Boxplots showing the F1-scores for cell type classification on the bone marrow dataset, comparing predictions by the four methods. N=16,325 image patches. **(c)** Boxplots showing the F1-scores for cell type classification on the sarcoma CODEX dataset, comparing predictions by the four methods. N=45,982 image patches. **(d)** A representative example of cell type clustering from the bone marrow dataset. Panels display, from left to right: the input H&E image patch, the ground-truth matched CODEX image with clustering, and the clustering results from the four methods. FMI scores are shown for each clustering result. **(e)** A representative example of cell type annotation from the bone marrow dataset using ground-truth annotation. The F1 scores for annotation accuracy are shown next to the method’s name. **(f)** A representative example of cell type annotation from the Sarcoma dataset using ground-truth annotation. The F1 scores for annotation accuracy are displayed next to the method’s name. P-values were calculated using a one-sided t-test.

Qualitative examples in Fig. 4d and 4e visually corroborate our findings using FMI and macro F1 metrics. The cell type maps generated by TissueCraftAI showed a spatial arrangement of cell populations that more closely resembles the ground truth maps. For example, using bone marrow H&E images, TissueCraftAI’s output (FMI= 0.35, Fig. 4d; F1=0.29, Fig. 4e) correctly delineated the spatial distributions of different cell types, a structure that is largely lost in the predictions by Pix2Pix-Turbo and HistoPlexer (Fig. 4e). Similar performance can be seen using sarcoma H&E images. For instance, TissueCraftAI correctly annotated rare endothelial cells (6% of all sarcoma cells), which were missed by the other two methods (Fig. 4f, Extended Data Fig. 4c). In sum, these results demonstrate the practical value of TissueCraftAI in enabling detailed, cellular-level analysis of the tissue microenvironment from standard histology images.

### High-Fidelity Prediction of Spatial Transcriptomics from H&E Images

We evaluated TissueCraftAI’s ability to predict spatial RNA expression directly from H&E images using the PRISM-12M dataset, comparing its performance to that of Pix2Pix-Turbo as a baseline method. Across 112 genes in the test data set, TissueCraftAI achieved a median F1 score of 0.24 compared to a 0.15 by Pix2Pix-Turbo (p = 0.0002). Fig 5b. shows the F1 scores for a set of representative genes. For a representative H&E image, TissueCraftAI accurately predicted the spatial locations of the *KRT16* spots, which closely matched the ground truth distribution. This accuracy is further demonstrated by a high Kernel Density Estimate (KDE) similarity of 0.50. In contrast, Pix2Pix-Turbo failed to predict the majority of true spots, resulting in a sparse and inaccurate density map, with a negative KDE similarity score of -0.05. Overall, our quantitative evaluation shows that TissueCraftAI significantly outperformed the baseline method across all metrics.

**Figure 5.**
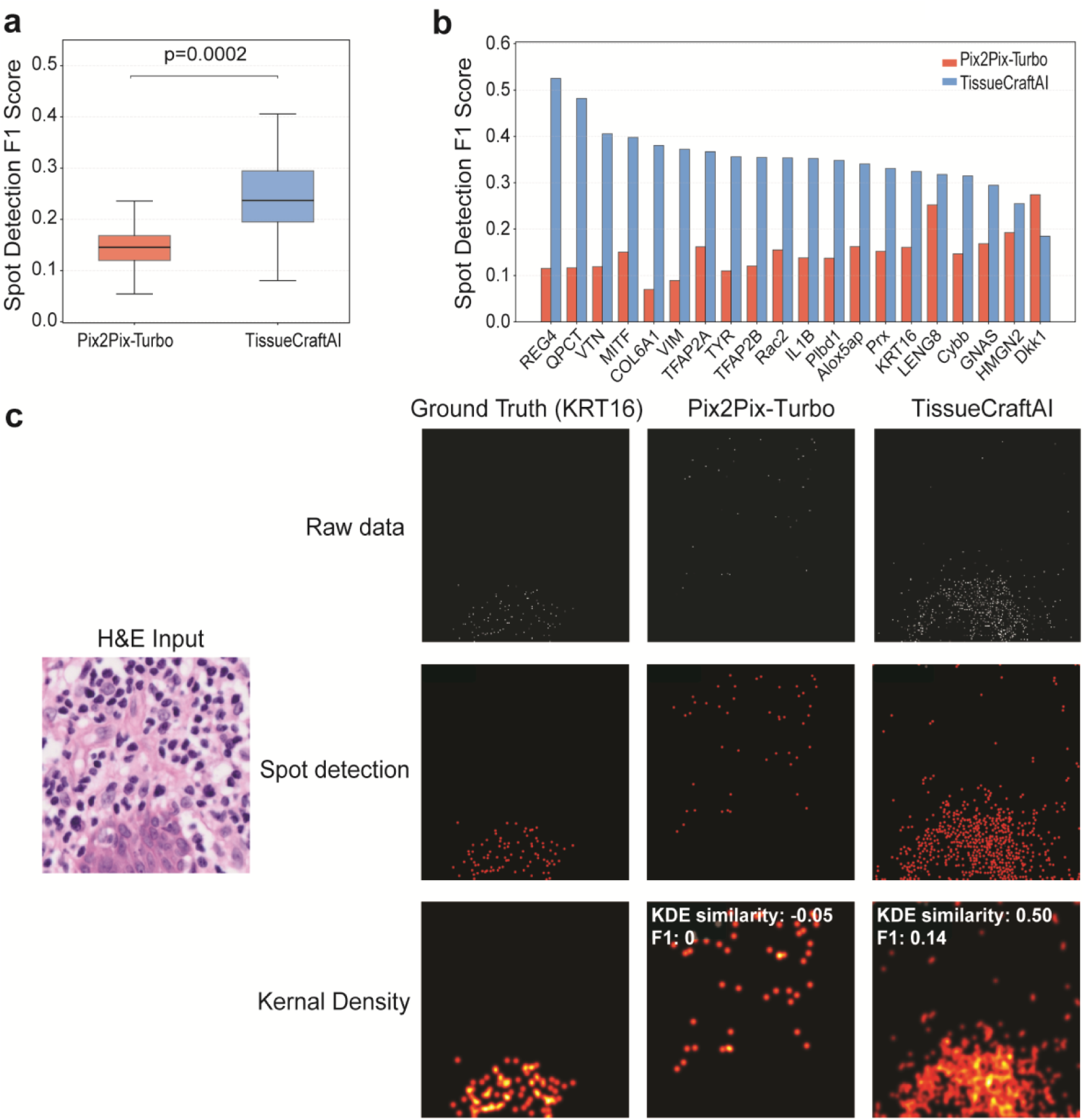
Prediction of transcript expression using histology images. (**a**) Box plot showing the F1 score distribution for RNA spot detection for each gene by Pix2Pix-Turbo and TissueCraftAI. The p-value from a one-sided t-test is shown. n=112 genes. **(b)** Bar plot of F1 scores for RNA spot detection for 20 representative genes. **(c)** Qualitative comparison of predicted spatial transcript distributions for a representative H&E image patch. From left to right: H&E input, ground truth transcript locations, predictions by Pix2Pix-Turbo, and predictions by TissueCraftAI. The bottom row displays kernel density estimations (KDE) of the predicted spots and corresponding F1 scores.

**Figure 6.**
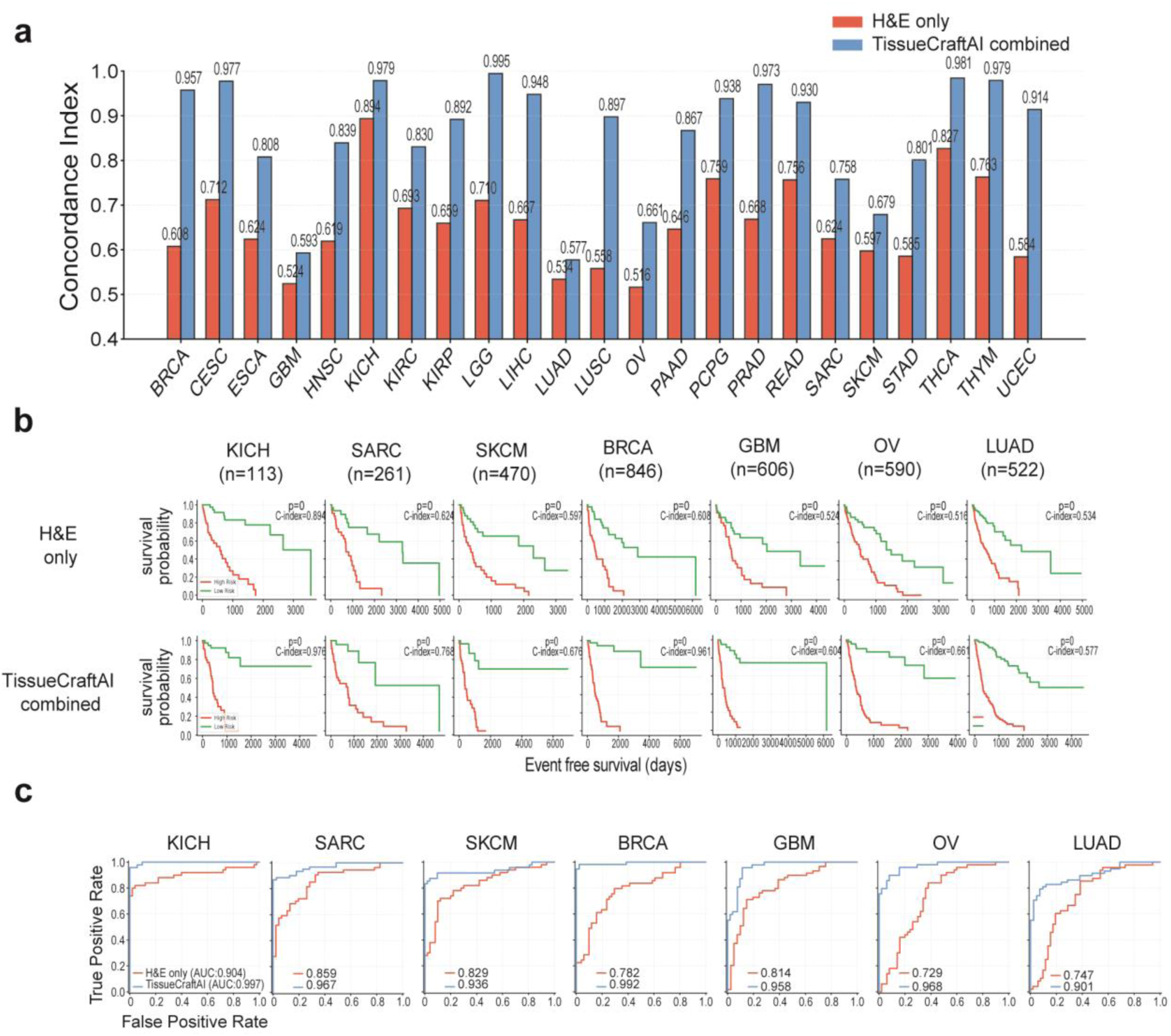
Enhanced prognostic stratification by integrating imputed proteomics with H&E image. **(a)** Concordance index (Harrell’s C-index) of prognostic models trained with H&E-only features versus models that combine H&E features with TissueCraftAI-inferred molecular maps across twenty-three TCGA cancer types: BRCA (Breast Invasive Carcinoma, n=846), CESC (Cervical Squamous Cell Carcinoma and Endocervical Adenocarcinoma, n=307), ESCA (Esophageal Carcinoma, n=185), GBM (Glioblastoma Multiforme, n=606), HNSC (Head and Neck Squamous Cell Carcinoma, n=523), KICH (Kidney Chromophobe, n=113), KIRC (Kidney Renal Clear Cell Carcinoma, n=535), KIRP (Kidney Renal Papillary Cell Carcinoma, n=291), LGG (Brain Lower Grade Glioma, n=516), LIHC (Liver Hepatocellular Carcinoma, n=377), LUAD (Lung Adenocarcinoma, n=522), LUSC (Lung Squamous Cell Carcinoma, n=504), OV (Ovarian Serous Cystadenocarcinoma, n=590), PAAD (Pancreatic Adenocarcinoma, n=185), PCPG (Pheochromocytoma and Paraganglioma, n=179), PRAD (Prostate Adenocarcinoma, n=500), READ (Rectum Adenocarcinoma, n=171), SARC (Sarcoma, n=261), SKCM (Skin Cutaneous Melanoma, n=470), STAD (Stomach Adenocarcinoma, n=443), THCA (Thyroid Carcinoma, n=507), THYM (Thymoma, n=124), and UCEC (Uterine Corpus Endometrial Carcinoma, n=560). Values above bars indicate cohort-level C-index. **(b)** Kaplan-Meier plots showing risk stratification based on model-derived risk scores; patients were divided at the median into high- and low-risk groups. Cohort size, C-index, and log-rank P value are shown. Top row: H&E-only features; bottom row: combined H&E + TissueCraftAI features. **(c)** Receiver operating characteristics (ROC) curves comparing event discrimination for H&E-only versus combined models; Areas Under the Curve (AUC) are shown.

### Zero-shot Application of TissueCraftAI Enhances Cancer Survival Prediction

To evaluate the clinical utility of TissueCraftAI, we investigated whether the synthesized spatial omics data it generates can improve patient survival prediction when used in conjunction with H&E images alone. We conducted a survival analysis across twenty-three cancer types from The Cancer Genome Atlas (TCGA) project (Fig. 6a). We comparison involved a Cox proportional-hazards survival model trained solely on H&E image features (”H&E only”) and a combined model that incorporated both H&E features and spatial omics features generated by TissueCraftAI (”TissueCraftAI combined”).

Our analysis workflow began with extracting high-quality 512x512 H&E image patches from TCGA whole-slide images. We then applied TissueCraftAI in a zero-shot manner to generate a corresponding set of synthesized spatial proteomics maps for each H&E image patch. Finally, we trained and compared two Cox proportional-hazards models. The “H&E only” model used deep features derived exclusively from the H&E patches using a ResNet50^19^ convolutional neural network, which were aggregated at the patient level. The “TissueCraftAI combined” model concatenated H&E features with those from the synthesized spatial proteomics maps at the patch level before aggregating at the patient-level. This approach created a richer, multimodal feature set for the survival prediction task.

The addition of TissueCraftAI-generated data significantly enhanced the predictive accuracy of survival models in across twenty-three cancer types. This was quantified by the Concordance Index (C-index)^20^, where the combined model outperformed the H&E-only model in all cohorts (Fig. 6a). The most notable improvements were seen in breast cancer (BRCA), where the C-index increased from 0.608 to 0.957; lung squamous cell carcinoma (LUSC), which improved from 0.556 to 0.873; and ovarian cancer (OV), with an increase from 0.536 to 0.856.

Kaplan-Meier curves further demonstrated the enhanced risk stratification abilities of TissueCraftAI (Fig. 6b, Extended Data Fig. 7). The combined model consistently achieved a more significant separation between high-risk and low-risk patient groups. For example, in the BRCA cohort, the statistical significance of the separation improved substantially with a p-value of 0.019 for the H&E-only model compared to 1.3e-4 for the combined model. Additional evidence of improved predictive accuracy is provided by the Receiver Operating Characteristic (ROC) curves (Fig. 6c, Extended Data Fig. 8), which show that the Area Under the Curve (AUC) was higher for the combined model across all cohorts. These results strongly suggest that the molecular information generated by TissueCraftAI provide valuable prognostic information that is not fully captured by tissue morphology alone.

## Discussion

In this study, we introduce TissueCraftAI, a text-guided generative AI framework that accurately predicts multi-modal spatial omics data from standard histology images. Our results demonstrate that TissueCraftAI outperforms existing state-of-the-art methods in generating high-fidelity spatial maps for both proteins and RNA transcripts. Additionally, we demonstrated that the *in silico* data generated by our model is useful for various downstream biological analyses, including improving the accuracy of spatial cell type annotation and significantly enhancing the prognostic power of clinical survival models. The development of this tool was made possible by the creation of PRISM-12M, a large-scale, spatially registered dataset that serves as a valuable resource for the computational pathology community.

TissueCraftAI introduces two major innovations. First, it features an integrated architecture that combines stable diffusion with ControlNet, allowing the model to accurately anchor the generation of molecular maps to the specific morphological context of input histology slides. The second innovation is its flexible, language-conditioned design, which represents a significant shift from the fixed, marker-specific models that have previously limited computational pathology. By framing molecular prediction as a text-guided generative task, TissueCraftAI allows for an open-ended exploration of tissue. Users can dynamically query the tissue microenvironment for any molecular target without requiring computational retraining or additional tissue sections. This natural language interface also positions TissueCraftAI as a foundational element for emerging biomedical AI agents.

While our results are promising, it is important to recognize the limitations of this study. The predictions made by TissueCraftAI are, by their nature, *in silico* and require experimental validation in specific research or clinical contexts. Although the model has been trained on a diverse dataset, its performance may vary for tissue types, disease states, or molecular markers that are not well-represented in PRISM-12M. Therefore, its ability to generalize will need continuous assessment as more data becomes available. Future work will focus on expanding the PRISM-12M resource to include a broader range of tissue types and omics modalities, such as spatial metabolomics and epigenomics.

In conclusion, TissueCraftAI represents a major advancement in computational pathology. It offers a flexible and accurate connection between routine histology and high-dimensional spatial omics, establishing a powerful new approach to tissue analysis. This work not only presents an innovative computational tool but also delivers a foundational dataset that will encourage further advancements, ultimately aiming to transform the study of tissue microenvironment.

## Online Methods

### Development of the PRISM-12M dataset

#### Spatial proteomics data processing and quality control

Spatial omics images underwent rigorous quality control procedures to ensure the curation of high-quality data for downstream analysis. Each image underwent manual inspection by trained researchers to identify and exclude samples with significant technical artifacts, including tissue folding, air bubbles, or staining irregularities. To remove outlier pixels, we applied an intensity threshold filter that excluded pixels with values exceeding the 99th percentile of the intensity distribution within each image. This conservative threshold was chosen to preserve biological heterogeneity while eliminating extreme outliers that could arise from autofluorescence, dust particles, or imaging artifacts.

#### Spatial transcriptomics data processing and quality control

For each sample image, we generated a processed image with the dimension of (X, Y, n_RNAs), where X and Y correspond to the spatial coordinates of a pixel, and n_RNAs is the number of RNA species. This representation preserves the spatial context while enabling analysis using a convolutional neural network. Expression values were normalized using log1p transformation followed by min-max scaling to the range [0, 255] for compatibility with standard image formats. Due to the presence of RNA molecules with extremely low signals, we developed a strategy to select image patches with robust RNA signals as high-quality data. First, for each RNA species, we calculated its total signal intensity across all image patches in a sample. Only image patches with a total signal for the given RNA intensity above the 75^th^ percentile were retained.

#### Image registration

To facilitate paired learning between imaging modalities, we aligned H&E histology images with spatial omics images using the wsireg toolkit (https://github.com/NHPatterson/wsireg). Wsireg performs image registration with an initial affine transformation based on tissue boundaries, followed by elastic deformation to adjust for local tissue distortions.

#### Tissue cellular content detection and image patch extraction

Registered images were systematically divided into non-overlapping patches measuring 512 × 512 pixels, which corresponds to approximately 256 × 256 μm at a resolution of 0.5 μm per pixel. This patch size strikes a balance between computational efficiency and providing sufficient spatial context for analyzing cellular structures. To eliminate background regions, we applied an intensity-based filter to the H&E patches, retaining only those with a mean RGB intensity of less than 220 (on a scale of 0-255). This threshold effectively removes white background areas while preserving patches that contain cellular material, including both densely cellular and stromal regions. The threshold was determined empirically by analyzing the intensity distribution across 1,000 manually annotated tissue and background patches.

#### Task formulation and text prompts

We formulated the multi-modal learning task as a conditional image generation problem with natural language conditioning. For predicting protein and RNA expression, we used structured text prompts formatted as: “predict [protein/gene name] protein/RNA expression”, which explicitly specifies the target molecular modality and identity. For the DAPI-to-H&E translation task, we used the prompt “predict corresponding H&E image”, framing the nuclear stain as input for predicting the histological appearance. These prompts enable flexible task conditioning within a unified framework and can be adapted to additional modalities or prediction targets.

#### Dataset partitioning for model training and testing

To ensure robust model evaluation and prevent data leakage, we partitioned the PRISM-12M dataset at the donor level using stratified random sampling, resulting in an 80:10:10 split for the training, validation, and test sets, respectively. By using donor-level splitting, we ensured that all patches from a specific donor were contained within a single subset. This approach helps to avoid overly optimistic performance estimates that could result from spatial autocorrelation within slides. We performed stratification based on tissue type and disease status to maintain balanced representation across the subsets. The validation set was used for hyperparameter tuning and model selection, while the test set was kept completely separate until the final evaluation.

### TissueCraftAI

#### Stable diffusion module

Stable Diffusion is a prominent generative model, specifically a type of Latent Diffusion Model (LDM), that is highly effective for text-to-image synthesis and other conditional generation tasks^11,12,14,21^. It operates based on the principles of diffusion probabilistic models, which involve a forward process that gradually adds Gaussian noise to data and a learned reverse process that denoises a random noise sample back into a coherent data point. A key innovation of Stable Diffusion is that it performs this diffusion process not in the high-dimensional pixel space, but within a lower-dimensional latent space learned by a powerful Variational Autoencoder (VAE). This significantly reduces computational requirements while maintaining high fidelity, a critical advantage when generating high-resolution histology or spatial omics images. The reverse denoising process is typically parameterized by a time-conditioned U-Net architecture. This U-Net iteratively predicts and removes noise from the latent variable *z*_*t*_ at timestep *t*, crucially guided by conditioning information *c* (e.g., image and text embeddings from the encoders) injected via mechanisms such as cross-attention. The primary objective during training is often to predict the noise *ε* added at a given step. This prediction, made by the U-Net model *ε*_*θ*_ with parameters *θ*, can be represented as:

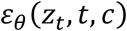

Here, *ε*_*θ*_ aims to approximate the actual noise ε added to reach the noisy latent state *z*_*t*_ at timestep *t*, given the conditioning context *c*. By iteratively applying this learned denoising function, the model transforms an initial random latent vector into a structured latent representation reflecting the condition *c*, which is finally decoded by the VAE’s decoder into the generated biomedical image.

#### ControlNet module

ControlNe^13,15^ introduces a method for adding fine-grained spatial conditioning to large, pre-trained text-to-image diffusion models, such as Stable Diffusion, without compromising their original capabilities learned from vast datasets. It achieves this by creating a trainable copy of the diffusion model’s encoder blocks (typically within the U-Net backbone) while keeping the original weights frozen. This trainable branch extracts multi-scale morphological features from the histology input. These features are passed through “zero convolution” layers (1x1 convolutions initialized with zeros) and added to the feature maps of the frozen denoising U-Net. This ensures that the morphological guidance is injected into the iterative denoising process without disrupting the semantic features derived from the text encoder. The effective operation within a specific block *i* of the U-Net can be conceptually represented as:

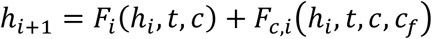

Here, *h*_*i*_ is the input feature map to block *i*, *F*_*i*_ represents the operation of the original frozen block, and *F*_*i*,*i*_ represents the operation of the trainable ControlNet copy for that block, which incorporates the additional spatial condition *c*_*f*_. *h*_*i*+1_ is the resulting feature map passed to the next layer. This additive mechanism enables ControlNet to provide precise spatial guidance – whether from morphological features in tissue sections or molecular distribution patterns – while leveraging the foundational model’s generative capacity.

### Running of Pix2Pix-Turbo

Pix2pix-turbo^16^ (https://github.com/GaParmar/img2img-turbo) adapts Stable Diffusion Turbo for single-step image-to-image translation by fine-tuning the U-Net and VAE with Low-Rank Adaptors (LoRAs). The method encodes input images through a modified VAE that captures skip connections, then handles the latent variables with a LoRA-adapted UNet that incorporates text conditioning. The output is decoded using the preserved skip connections for enhanced reconstruction. The Training uses a multi-loss objective that combines L2 loss (λ=1.0), Learned Perceptual Image Patch Similarity (LPIPS) loss (λ=5.0), Contrastive Language-Image Pretraining (CLIP) loss (λ=5.0), and adversarial losses (λ=0.5) to produce high-quality outputs. A vision-aided GAN discriminator using CLIP features and multilevel sigmoid loss is also used. The model supports both deterministic inference and stochastic modes with controllable noise blending, enabling efficient training with paired data while preserving the semantic understanding of pre-trained diffusion models in a single forward pass. For our experiments, we trained the model on four GPUs with an effective batch size of eight (two per GPU) at a resolution of 512×512 pixels for 500,000 steps. We used the AdamW optimizer with a learning rate of 5×10⁻⁶, β₁=0.9, β₂=0.999, and a weight decay of 0.01. LoRA ranks were set to eight for the U-Net and four for the VAE. We enabled memory-efficient attention via xformers and applied gradient clipping with a maximum norm of 1.0.

### Running of HistoPlexer

We used HistoPlexer^9^ (https://github.com/ratschlab/HistoPlexer) with its default configuration settings as specified in the official implementation. HistoPlexer uses conditional generative adversarial networks (cGANs) with custom loss functions designed to mitigate slice-to-slice variations and preserve spatial protein correlations. Training was conducted with a batch size of 16, using the Adam optimizer with a learning rate of 1e-4 and a learning rate scheduler with a decay rate of 0.8 and patience of 15 epochs. The model was trained for a maximum of 100 epochs with early stopping enabled to prevent overfitting. The method processes high-resolution histopathology images by default (use_high_res: True) and generates spatially resolved protein expression. The loss function incorporates Gaussian Process (GP) regularization with a weight of 5.0 (w_GP: 5.0) and Asymmetric Spatial Prior (ASP) with a weight of 1.0 (w_ASP: 1.0). Training utilized 8 worker processes for data loading (num_workers: 8) and employed a random seed of 96 for reproducibility. For inference, we used the trained model checkpoints to generate protein predictions directly from H&E images, maintaining the same preprocessing pipeline and architectural specifications as used during training.

### Metrics and procedures for evaluating the quality of synthetic images

To quantitatively evaluate the quality and diversity of the generated images, we used the Fréchet Inception Distance (FID) score^17^. The FID score measures the similarity between two sets of images by comparing the statistics of their feature representations, as extracted from an intermediate layer of a pre-trained InceptionV3 network. These feature distributions, for both the real images (x) and the generated images (g), are modeled as multivariate Gaussian distributions. The distance between these two distributions is then calculated using the Fréchet distance, defined by the following formula:

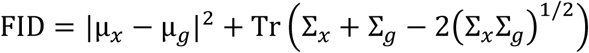

In this equation, μ_*x*_ and μ_*g*_ represent the means of the feature vectors for the real and generated images, respectively, while Σ_*x*_ and Σ_*g*_ are their corresponding covariance matrices. The term Tr denotes the trace of the matrix. A lower FID score indicates that the distribution of generated images is closer to the distribution of real images, suggesting higher quality and greater diversity in the generated samples.

Alongside FID, we used the Pearson Correlation Coefficient (PCC) to measure the linear correlation between the pixel intensity values of the generated image (X) and the ground truth image (Y). PCC is calculated as:

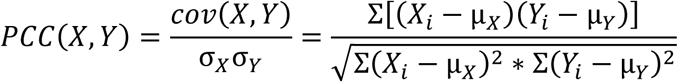

where *X*_*i*_ and *Y*_*i*_ are the intensity values of the i-th pixel, *σ*_*X*_ and *σ*_*Y*_ are the respective mean intensities, *σ*_*X*_ and *σ*_*Y*_ are the standard deviations, and the summation is over all pixels. Together, FID and PCC provide complementary assessments of structural similarity and intensity correlation for evaluating the quality of the synthesized images.

### RNA spot detection algorithm

We developed a pipeline for identifying individual RNA molecules as discrete punctate signals in spatial transcriptomics images. The pipeline employs adaptive local thresholding to account for spatial variations in background intensity. For each input image *I*, we first normalized the pixel intensities to the range of [0,1] and applied Gaussian smoothing with σ = 1.0 to reduce noise. The binary mask B was then computed using the following formula:

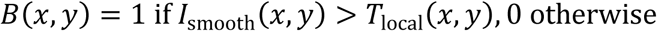

where *T*_*local*_ is the local threshold computed using a Gaussian-weighted window of size 15×15 pixels. Connected components with an area of less than 3 pixels were removed as noise.

### Spot matching and F1 score calculation

To evaluate the detection accuracy, we used a nearest-neighbor matching algorithm to compare the predicted and ground-truth centroids of RNA spots. For predicted spots *P* = {*p*_1_, *p*_2_, …, *p*_*n*_} and ground-truth spots *G* = {*g*_1_, *g*_2_, …, *g*_*j*_}, we computed the Euclidean distance matrix D, where *D* = ‖*p* − *g* ‖ . A predicted spot *p*_i_ was considered a true positive if its nearest ground truth spot *g*_*j*_ satisfied *D*_*ij*_ ≤ τ (where τ = 10 pixels) and *g*_*j*_ had not been previously matched. The precision, recall, and F1 score are calculated as:

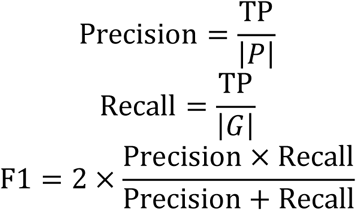

where TP is the number of true positive matches.

### Kernel density estimation for spatial distribution analysis

To capture larger spatial distribution patterns of RNA transcripts beyond individual spots, we implemented a kernel density estimation (KDE) approach. For a set of detected spots, *D* = {(*x*_1_, *y*_1_), (*x*_2_, *y*_2_), …, (*x*_*n*_, *y*_*n*_)}, the KDE image K was computed as:

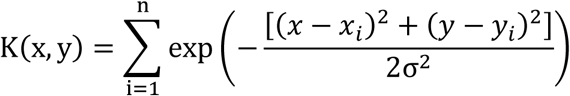

where *σ* is the bandwidth parameter (default *σ* = 5 pixels). This transformed the discrete spot pattern into a continuous density map that captures both local clustering and global distribution patterns.

### KDE similarity metric

To quantify the similarity between predicted and ground truth spatial distributions, we computed the Pearson Correlation Coefficient between their respective KDE representations:

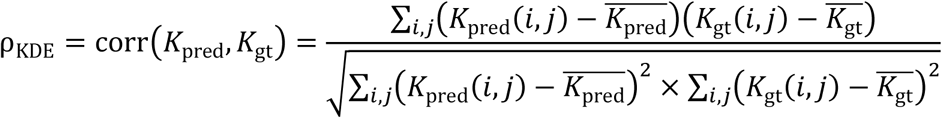

The KDE similarity is robust to small positional errors in individual spot detection while capturing the overall spatial organization of RNA transcripts.

### Clustering and cell type annotation of synthetic TissueCraftAI images from H&E images

The workflow began with spatial proteomics images from the Ground Truth, as well as synthetic images generated by TissueCraftAI and Pix2Pix-Turbo. We first applied the Mesmer algorithm for cell segmentation of these images, generating a cell mask for each sample. Using these masks, we quantified the mean expression intensity for each protein marker within each segmented cell boundary. This procedure yielded a cell-by-protein expression feature matrix, which served as the input for downstream analysis. For H&E images, RGB color-based features were extracted for each cell defined by the same cell segmentation mask from the matched spatial omics images.

After cell segmentation and expression quantification, we applied a uniform analysis pipeline for each of the four types of images (Ground Truth, H&E, Pix2Pix-Turbo, and TissueCraftAI). The feature matrix for each cell was first normalized to a total count of 10,000, followed by a logarithmic transformation. Next, we performed Principal Component Analysis (PCA) on the scaled feature matrix to reduce dimensionality. The number of principal components was chosen adaptively based on the feature space (up to 50 for protein data, 2 for low-dimensional H&E data). Using this PCA embedding, we constructed a k-nearest neighbor graph (k=20) and subsequently performed Leiden clustering (resolution=0.5) to partition cells into distinct clusters. For visualization, a Uniform Manifold Approximation and Projection (UMAP) embedding was computed from the PCA representation.

An automated cell type annotation strategy was employed to assign cell type identities. For protein-based modalities (Ground Truth, TissueCraftAI, Pix2Pix-Turbo), we scored the mean expression of canonical protein markers for each cell type. Each cell was assigned an identity based on its highest-scoring marker set. For the H&E modality, the RGB color density values of each cell were normalized and log-transformed. Leiden clustering was then applied to identify cell clusters. Clusters were mapped to cell type labels (tumor, epithelial, immune) using the Hungarian algorithm to find the optimal matching that maximizes overlap with ground truth annotations.

The cell type annotation accuracy using each data modality was evaluated against the ground truth annotations. We used two categories of metrics: 1) Clustering Similarity: To determine the similarity of two clusterings, we calculated the Fowlkes-Mallows Index (FMI)^18^. 2) Cell Type Classification Accuracy: We computed accuracy, precision, recall, and the F1-Score. To ensure a fair comparison across different methods, we first found the optimal assignment between the test and reference cell type labels by applying the Hungarian algorithm to the confusion matrix.

Finally, aggregated metrics across all samples were visualized, and a t-test was used to calculate statistical significance between TissueCraftAI and the other methods.

### Patient survival analysis

Our survival prediction workflow consisted of three sequential steps: extraction of H&E image patches from whole-slide images (WSIs), generation of synthesized spatial proteomics maps using TissueCraftAI, and training and evaluation of survival models.

#### H&E image patch extraction and processing

To create a managed dataset of high-resolution image patches from Whole Slide Images (WSIs) of tumor tissues from The Cancer Genome Atlas (TCGA) project^22^, we developed a custom processing pipeline. First, we generated a low-resolution thumbnail for each WSI to facilitate efficient tissue detection. We converted this thumbnail into the HSV color space and applied a color saturation threshold of 0.15 to create a binary tissue mask, which effectively isolated tissue regions from the background of the empty slide. Next, we sampled up to 100 non-overlapping 512×512-pixel patches exclusively from the masked areas of the high-resolution WSI. This number was chosen to standardize the dataset, as it represents the approximate number of quality patches typically obtainable from a single slide. To ensure that each patch contained significant tissue content, we conducted a quality control check, discarding any patch with a mean pixel intensity above 220 (on a 0-255 scale). This step helped filter out regions with minimal or no tissue. The final extracted H&E patches were saved in a structured directory organized by patient. Additionally, we generated a corresponding metadata file to log the precise coordinates of each patch within its original WSI, ensuring full traceability.

#### In silico spatial proteomics image generation

In the second stage, we used the directories of H&E patches as input to our TissueCraftAI model. Through a deep learning inference process, the model generated a corresponding set of spatial protein expression maps for each H&E patch. The output was a parallel directory of “synthesized protein maps” where each input H&E patch was associated with multiple grayscale images. These images represented the predicted spatial expression of individual protein markers.

#### Survival model training and evaluation

In the final stage, we used both H&E patches and their corresponding synthesized protein maps to develop and compare survival prediction models. We ran two distinct feature extraction pipelines for each patient. The H&E-only model focused on extracting deep features exclusively from the H&E patches using the pretrained convolutional neural network (CNN), ResNet50.

These features were then aggregated into a single feature vector for each patient, comprising the mean, standard deviation, maximum, and minimum values. The multimodal model extracted features from both the H&E patches and their associated synthesized protein images. For each patch, the H&E features were concatenated with the aggregated features from the corresponding synthesized protein channels, resulting in a final patient-level feature vector.

We constructed and compared three multivariate Cox proportional hazards models with elastic net regularization to predict overall survival. The first model used a baseline set of clinical variables, including age, sex, and tumor stage, along with features extracted from H&E images. The second model replaced the H&E features with those generated by TissueCraftAI while retaining the clinical variables. The third model combined features from H&E images, TissueCraftAI, along with clinical data. To handle the high-dimensional image features, we first applied Principal Component Analysis (PCA) for dimensionality reduction, followed by standardization. We assessed model performance using the Concordance Index (C-index) with 5-fold stratified cross-validation. To visualize the prognostic capabilities, we stratified patients into high- and low-risk groups based on the median predicted risk score. We generated Kaplan-Meier survival curves and compared them using the log-rank test. Finally, we assessed the statistical significance of the improvement provided by the TissueCraftAI features over the baseline model using the likelihood ratio test.

## Data availability

Information for all datasets used in this study is provided in Supplemental Table 1.

## Code availability

TissueCraftAI model weights and code to extract slide embeddings will be released upon publication at github.

## Acknowledgments

The authors thank the Children’s Hospital of Philadelphia Research Information Services for providing computing support. This work was supported by the National Cancer Institute (NCI) Human Tumor Atlas Network grant under award #U2C CA233285 (K.T.) and the National Institutes of Health (NIH) Human Biomolecular Atlas Program grant under award #U54 HL165442 (K.T.).

## Author Contributions

M.P. and K.T. conceived and designed the study. M.P. and K.T. curated and annotated the PRISM 12M dataset. M.P. implemented the TissueCraftAI algorithm and developed the software package. M.P. and T.R. performed data analysis, X.W. and K.T. supervised the overall study. M.P. and K.T. wrote the manuscript with input from all authors.

## Competing interests

The authors declare no competing interests.

## Notes

### Competing Interest Statement

The authors have declared no competing interest.

